# Preexisting antibodies can protect against congenital cytomegalovirus infection in monkeys

**DOI:** 10.1101/127647

**Authors:** Cody S. Nelson, Diana Vera Cruz, Dollnovan Tran, Kristy M. Bialas, Lisa Stamper, Huali Wu, Margaret Gilbert, Robert Blair, Xavier Alvarez, Hannah Itell, Meng Chen, Ashlesha Deshpande, Flavia Chiuppesi, Felix Wussow, Don J. Diamond, Nathan Vandergrift, Mark R. Walter, Peter A. Barry, Michael Cohen-Wolkowiez, Katia Koelle, Amitinder Kaur, Sallie R. Permar

## Abstract

Human cytomegalovirus (HCMV) is the most common congenital infection and a known cause of microcephaly, sensorineural hearing loss, and cognitive impairment among newborns worldwide. Natural maternal HCMV immunity reduces the incidence of congenital infection, but does not prevent the disease altogether. We employed a nonhuman primate model of congenital CMV infection to investigate the ability of preexisting antibodies to protect against placental CMV transmission. Pregnant, CD4+ T cell-depleted, rhesus CMV (RhCMV)-seronegative rhesus monkeys were treated with either standardly-produced hyperimmune globulin (HIG) from RhCMV-seropositive macaques or dose-optimized, potently RhCMV-neutralizing HIG prior to intravenous challenge with an RhCMV swarm. HIG passive infusion provided complete protection against fetal loss in both groups, and the potently-neutralizing HIG additionally inhibited placental transmission of RhCMV. Our findings suggest that antibody alone at the time of primary infection can prevent congenital CMV and therefore could be a primary target of vaccines to eliminate this neonatal infection.

## Introduction

The current epidemic of Zika virus in the Americas has raised significant awareness of the societal and pediatric health impact of a congenitally-transmitted neuropathogen.^1^ Yet nearly 1 million infants worldwide (1 out of every 150 live births) are born each year with congenital human cytomegalovirus (HCMV) infection.^2,3^ Congenital HCMV, like congenital Zika virus, can cause microcephaly, hearing/vision loss, and abnormalities in nervous system development.^2,4^ And though HCMV accounts for more congenital disease than all 29 newborn conditions currently screened for in the United States combined,^5^ public knowledge and effective interventions are severely lacking.^6,7^ In recognition of this burden of disease, a congenital HCMV vaccine has remained a “Tier 1 priority” of the National Academy of Medicine for the past 15 years.^8^

Drawing upon standard vaccine strategies, the HCMV field has previously attempted to induce potent antiviral immunity by varying approaches including attenuation of live viruses,^9-11^ formulation of HCMV glycoprotein subunit vaccines,^12-14^ use of viral vectors for epitope expression,^15-17^ and delivery of single and bivalent DNA plasmid vaccines.^18,19^ While all vaccine platforms tested thus far ultimately failed to reach target endpoints in human clinical trials, the HCMV gB subunit vaccine demonstrated moderate (∼50%) efficacy at preventing primary HCMV infection, which is promising for future vaccine-development efforts.^13,14^ Additional retrospective human studies have reported that neutralizing antibodies targeting HCMV surface glycoproteins are correlated with reduced incidence of congenital virus transmission after primary maternal HCMV infection.^20-22^ Yet, a recent clinical trial failed to demonstrate a significant reduction in rates of congenital HCMV infection following passive infusion of hyperimmune globulin (HIG) to pregnant women with primary HCMV infection. Results from these past studies emphasize possible challenges for the development of an efficacious antibody-based vaccine.^23^

One major barrier the HCMV vaccine field has faced is the lack of a highly-translatable animal model of congenital virus transmission. For the past 50 years, vaccine efficacy studies that evaluate protection against congenital HCMV transmission have been reliant upon either (1) small-animal models using species-specific viruses with limited HCMV sequence homology or (2) costly and arduous human clinical trials. We recently reported on the first nonhuman primate (NHP) model of congenital CMV transmission following primary infection of rhesus monkey dams with rhesus CMV (RhCMV),^24^ which is genetically more similar to HCMV than guinea pig or murine CMV.^24-26^ Our study established that RhCMV could cross the placenta and cause congenital infection following intravenous (IV) inoculation of either immune competent or CD4+ T cell-depleted seronegative dams during the second trimester of pregnancy. The IV route of RhCMV inoculation was selected for this experimental congenital infection model over mucosal routes of exposure as it induced reproducible high levels of viremia and virus shedding in the urine and saliva of infected macaques, and hence best mirrored the prerequisite for systemic CMV replication prior to placental virus transmission in congenital CMV infection. In this study, we observed that CD4+ T cell-depleted dams frequently aborted their fetus following virus inoculation, exhibited higher plasma and amniotic fluid viral loads, and had delayed production of autologous neutralizing antibodies.

These results suggested that maternal humoral immunity may impact systemic and intrauterine RhCMV replication, thereby influencing the severity of congenital infection. As such, we sought to use this NHP model to investigate whether preexisting antibody alone, in the absence of virus-specific T cell responses, could protect against placental RhCMV transmission or reduce congenital RhCMV infection severity. The passive infusion of antibodies in this study was not intended to model a treatment strategy for the prevention of either maternal CMV acquisition or fetal virus transmission, but rather as means to understand the protection conferred by preexisting maternal antibody. These insights will inform rational development of a vaccine to elicit the most critical immune responses required to eliminate congenital HCMV disease.

## Results

### Hyperimmune globulin (HIG) production and study design

Two separate preparations of HIG were purified from plasma of RhCMV-seropositive rhesus monkey donors. The first “standard” preparation was obtained from donors with high levels of total RhCMV-specific IgG, while the second “high-potency” preparation was purified from plasma of donors exhibiting robust epithelial cell neutralizing IgG antibody titers (ID_50_>1:1000). The high-potency preparation had approximately 4-fold greater epithelial cell neutralization potency and slightly increased overall glycoprotein binding (Fig. 1B,C) than the standard preparation. All RhCMV-seronegative dams in our current and previous study were infused with a CD4+ T cell-depleting antibody at week 7 of gestation, as described.^24^ T cell phenotyping (Fig. S1, Table S1) revealed a decline in CD4+ T cells with changes in the CD8+ T cell population in response to primary RhCMV infection (Fig. S2). The dams were subsequently divided into 3 groups: control, standard, and high-potency. Each animal, including those in the control group (n=6, including 4 historical controls^24^) were IV inoculated with a swarm of fibroblast-tropic 180.92^27^ and epithelial-tropic UCD52/UCD59^28^ RhCMV viruses at week 8 of gestation (Fig. 1A). The standard group (n=3) was administered a single dose (100mg/kg) of the standard HIG preparation 1-hour prior to inoculation with the RhCMV variants (Fig. 1B), while the high-potency group (n=3) was given a dose-optimized regimen of the high-potency HIG preparation (Fig. S3) 1-hour prior to (150mg/kg) and 3 days following (100mg/kg) RhCMV inoculation (Fig. 1C, Fig. S4).

**Fig. 1.**
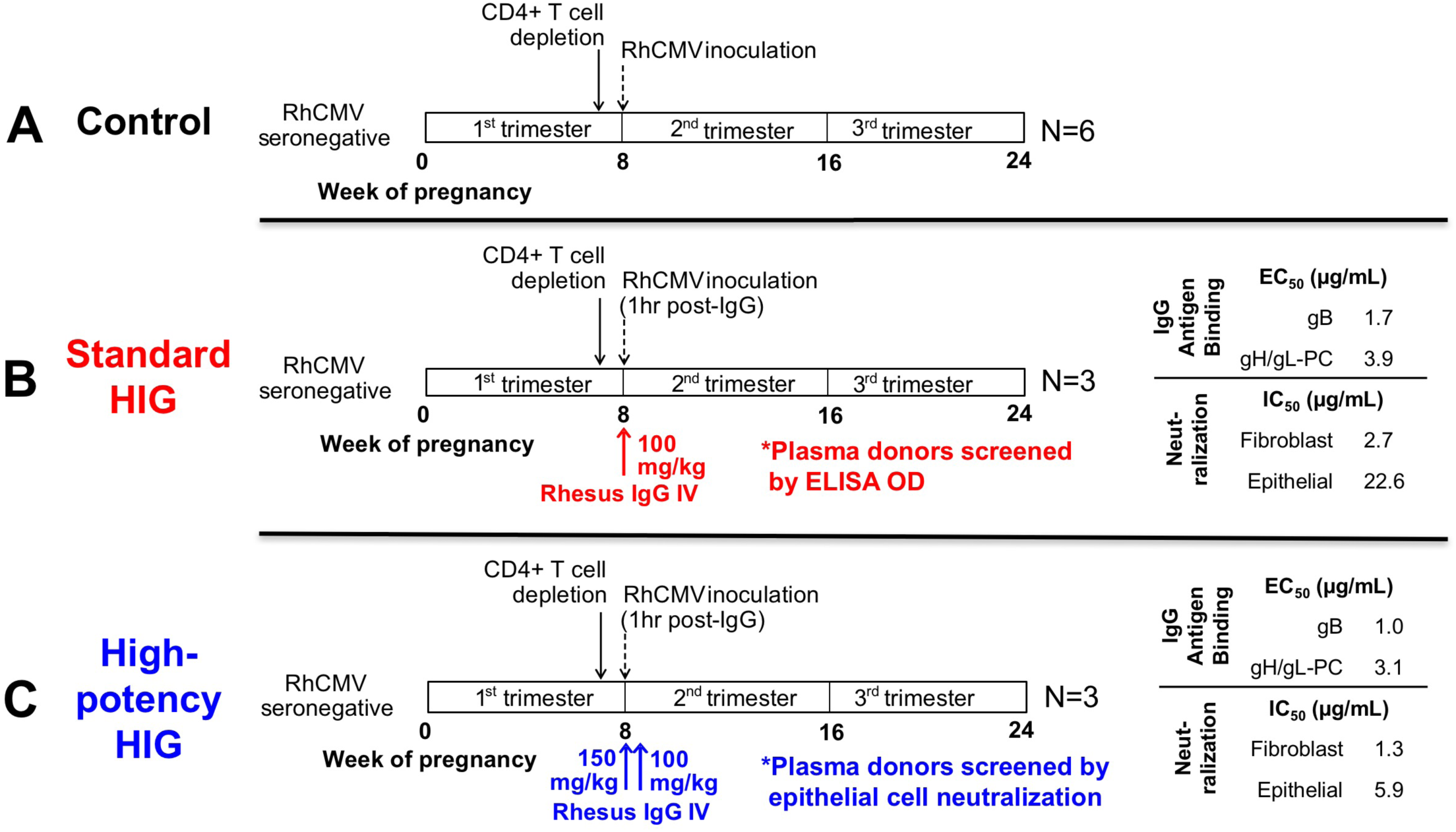
HIG preinfusion/RhCMV placental transmission study experimental design. (**A**) 6 RhCMV-seronegative rhesus monkey dams were CD4+ T cell depleted at week 7 of gestation then inoculated one week later with a swarm of RhCMV stocks (180.92, UCD52, UCD59). (**B**) 3 seronegative, CD4+ T cell-depleted dams received a single dose (100mg/kg) of a standard HIG preparation 1hr prior to RhCMV infection. (**C**) 3 seronegative, CD4+ T cell-depleted dams received a dose-optimized HIG regimen (150mg/kg prior to RhCMV infection +100mg/kg 3 days following infection) of a high neutralizing potency HIG preparation isolated from seropositive donors screened by epithelial cell neutralization. Quantified IgG glycoprotein complex-specific binding titers as well as fibroblast (180.92 virus) and epithelial (UCD52 virus) neutralization activity was assessed for each HIG preparation (**B,C**).

### Glycoprotein targets of RhCMV-neutralizing antibodies

To determine the RhCMV antibody specificity of the administered HIG, both standard (Fig. 2A) and high-potency HIG (Fig. 2B) were depleted for antibodies specific for RhCMV glycoprotein B (RhgB), RhCMV pentameric complex (RhPC), as well as RhgB/RhPC combined. Sufficient and selective depletion of glycoprotein-specific antibodies was confirmed by ELISA against the depleted epitope. We confirmed a >75% decrease in EC_50_ magnitude against depleted epitope and <20% change in EC_50_ against non-depleted epitope (Fig. 2C). For the standard HIG preparation, absorption with RhgB resulted in a 2.2-fold reduction in the neutralization potency of plasma (mock depleted IC_50_=32.3 vs. RhgB-depleted IC_50_=71.8), whereas absorption with RhPC resulted in a slightly higher 2.9-fold reduction in neutralization potency (RhPC-depleted IC_50_=94.6) (Fig. 2A,C). This same trend was observed for antigen depletion of the high-potency HIG, with a 2.9-fold reduction following RhgB depletion and a 3.3-fold reduction following RhPC depletion (Fig. 2B,C). Additionally, depletion of antibodies specific to *both* antigens reduced the neutralization potency of sera beyond that of depletion with either RhgB or RhPC alone (Fig. 2A-C). Altogether, these findings suggest that antibodies targeting both RhgB and RhPC contribute to overall RhCMV neutralization activity in roughly equivalent proportions. Furthermore, both RhgB and RhPC-specific antibodies were found to contribute to the RhCMV-neutralization titer in individual RhCMV-seropositive rhesus monkeys (Fig. S5).

**Fig.2.**
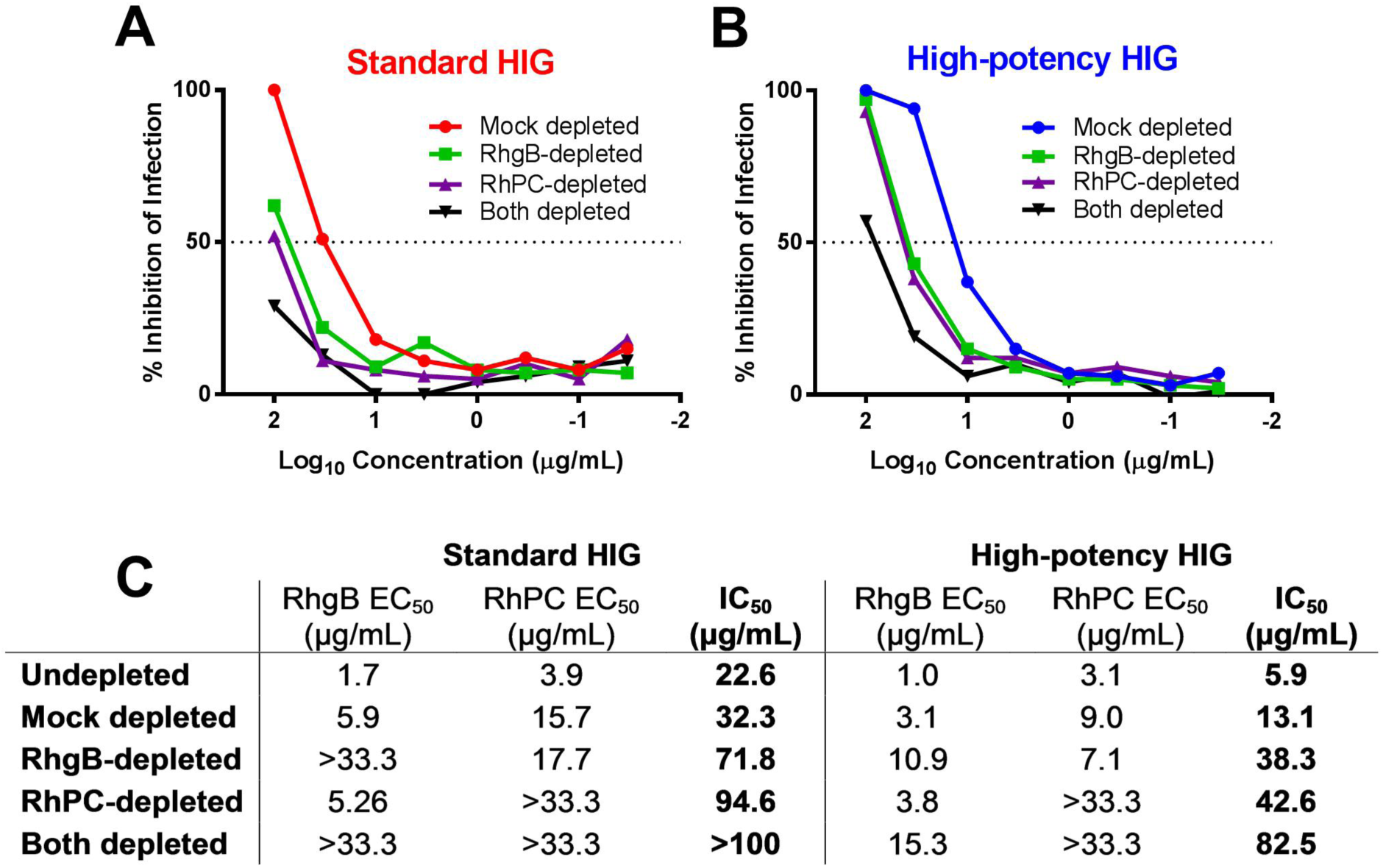
Both RhCMV gB and PC-specific antibodies contribute to neutralization activity of purified HIG preparations. Neutralization curves for standard HIG (**A**) and high-potency HIG (**B**) compare the relative neutralization potency of mock-depleted, RhgB-depleted, RhPC-depleted, and both RhgB/RhPC-depleted antibody preparations. (**C**) IC_50_ values for depleted antibodies measured in epithelial cells with UCD52 virus (starting concentration 100μg/mL). ELISA EC_50_ values (measured here in μg/mL RhCMV HIG equivalent) confirm specific depletion of antibodies for a given antigen (starting concentration 33.3μg/mL).

### Antibody kinetics following HIG infusion

Total RhCMV-binding IgM and IgG titers (Fig. 3A,B), RhgB and RhPC-specific IgG titers (Fig. 3C,D), and fibroblast/epithelial cell IgG neutralizing antibody titers (Fig. 3E,F), were measured following HIG infusion. Based on the known kinetics of antibody responses to primary viral infections, we attributed early detection of RhCMV-specific activity (within the first 10 days) to reflect passively-infused antibodies in the HIG preparations. Peak RhCMV-binding IgG as well as RhgB and RhPC-specific antibody titers in the high-potency HIG group surpassed those of the standard HIG group (Fig. 3B-D), while peak neutralization titers were similar between the two HIG groups (Fig. 3E-F). However, levels of IgG binding and antibody neutralization were sustained near peak levels for 10 days in the dose-optimized, high-potency HIG group compared to only 3 days in the standard HIG group (Fig. 3B-E). It is noteworthy that average binding IgG and epithelial cell neutralization titers in the high-potency HIG group surpassed those of chronically-infected rhesus monkeys (Fig. 3B,E, Fig. S5).^24^ The standard HIG group had the most robust natural host immune response, with a sustained elevation of RhCMV-specific IgM (Fig. 3A), as well as an exponential rise in binding IgG (Fig. 3B) and neutralization titers (Fig. 3D,E) that outpaced each of the other groups.

**Fig. 3.**
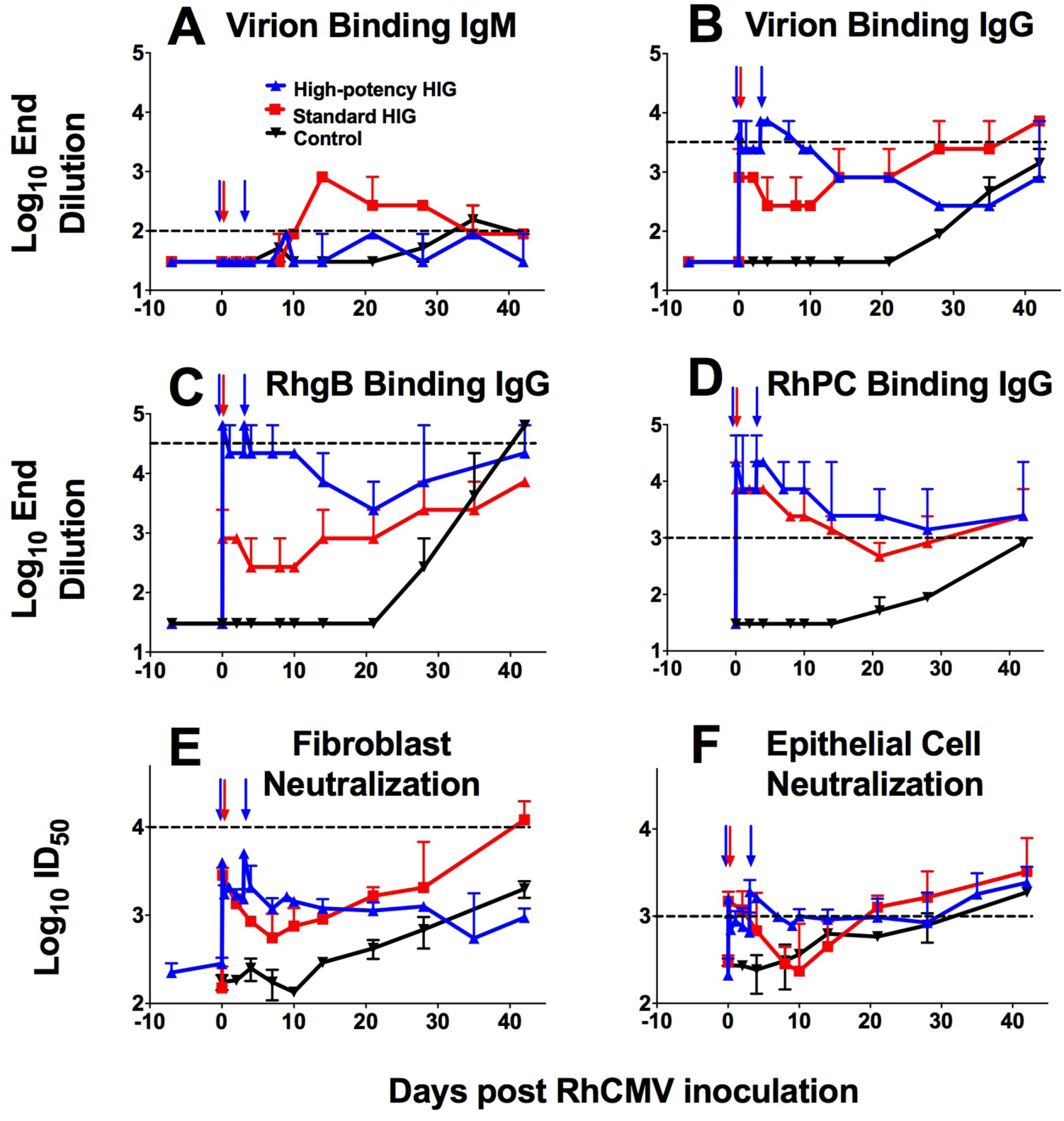
Plasma RhCMV IgM/IgG binding and neutralization following HIG infusion. Median whole UCD52 RhCMV virion IgM (**A**) and IgG (**B**) binding titers are shown for 2 control animals in black (excluding 4 historical controls because of sample limitation), the 3 animals infused with a single dose of standard HIG in red, and the 3 animals administered a dose-optimized regimen of high-potency HIG in blue. RhCMV glycoprotein B (RhgB) (**C**) and RhCMV pentameric complex (RhPC) (**D**) binding-IgG titers demonstrate higher glycoprotein binding in the high-potency HIG-infused group. Furthermore, median neutralization titers (ID_50_) were measured on fibroblast (180.92 virus) (**D**) and epithelial cells (UCD52 virus) (**E**). The time points of HIG infusion are denoted by colored arrows. Error bars represent the range of results between dams within the same treatment group. Horizontal dotted lines indicate average antibody binding and neutralization titers for chronically RhCMV-infected rhesus monkeys.

### RhCMV viral load, placental transmission, and shedding

Following RhCMV inoculation, both control and standard HIG group animals showed a rapid onset of viremia that peaked at 2 weeks post infection (Fig. 4A). In contrast, peak viremia in the high-potency HIG group was delayed until 4 weeks post infection (Fig. 4A), with a median value nearly two logs lower than the control group (Fig. 4E; p=0.047, corrected Wilcoxon exact test). RhCMV DNA copies in amniotic fluid were used as a marker for placental virus transmission because it is the gold-standard clinical test for congenital HCMV infection.^29^ All 6 control animals had detectable RhCMV DNA in amniotic fluid between 1 and 3 weeks following infection (Fig. 4B), indicating 100% placental transmission. Additionally, 2 of 3 animals in the standard HIG group had detectable congenital RhCMV infection (67% transmission) (Fig. 4B). However, none of the 3 animals in the high-potency HIG group had detectable RhCMV DNA in the amniotic fluid, suggesting complete inhibition of congenital RhCMV transmission (Fig. 4B), and indicating a potential dose-effect of HIG-mediated protection. It is possible that systemic viral load influenced placental transmission, as animals that transmitted the virus had the highest peak plasma viral loads (Fig. 4E) and peak plasma viral load correlated with initial amniotic fluid viral load (Fig. 4F; r=0.812, p=0.001, Spearman correlation). Finally, while there was no difference in the peak magnitude of viral shedding in maternal urine (Fig. 4C) or saliva (Fig. 4D) between treatment groups, there was delayed onset of shedding in both urine (Fig. 4G) and saliva (Fig. 4H) following high-potency HIG infusion (both p=0.047, corrected Wilcoxon exact test). Altogether, these findings suggest that the presence of antibodies can reduce systemic replication and block dissemination of the virus to other anatomic compartments.

**Fig. 4.**
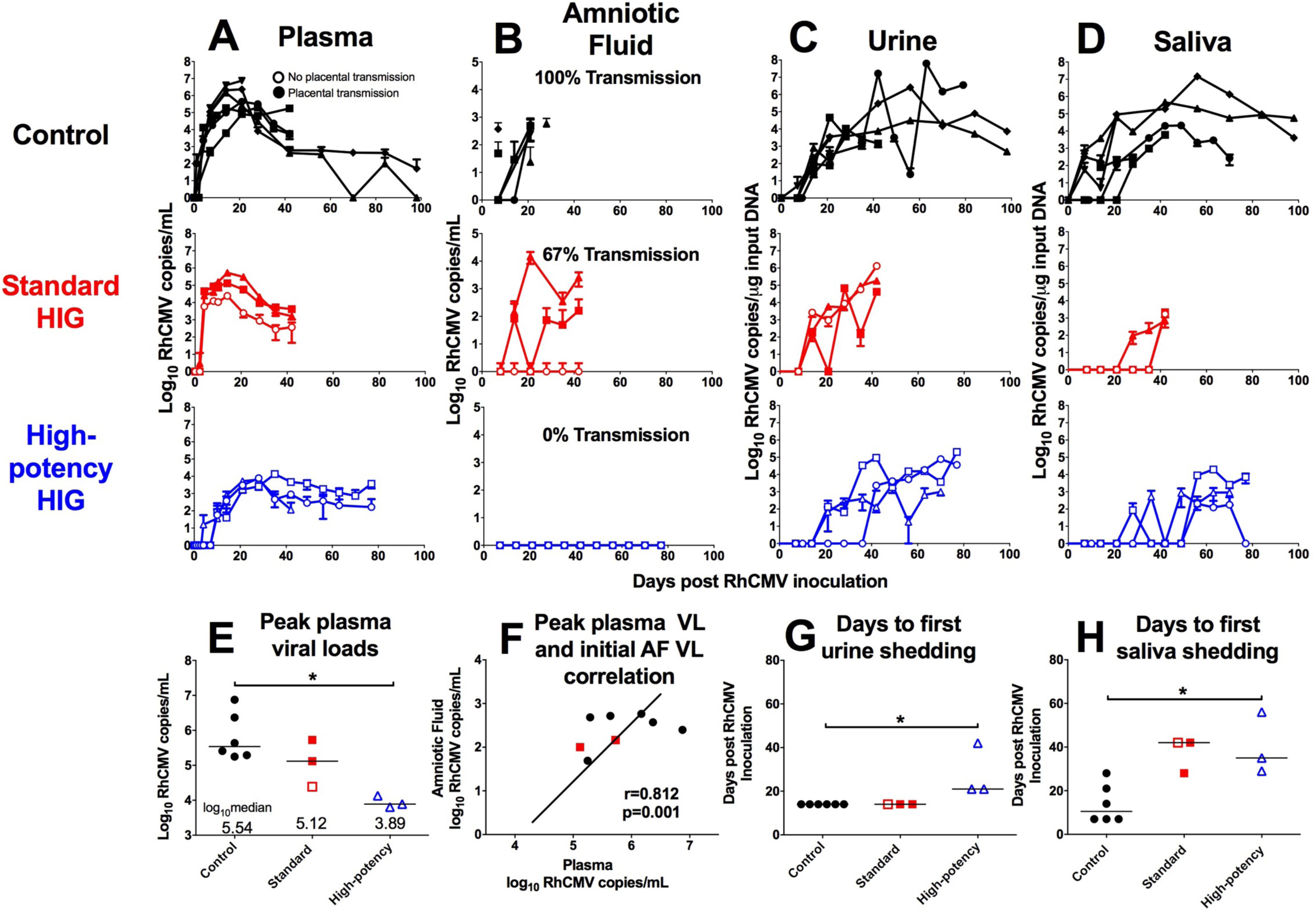
Infusion with high-potency HIG prior to RhCMV infection reduces plasma viral load, prevents placental virus transmission, and delays viral shedding. RhCMV copy number determined by qPCR is shown for control animals (n=6; black), the standard HIG group (n=3; red), and the high-potency HIG group (n=3; blue) in plasma (**A**),amniotic fluid (**B**), urine (**C**), and saliva (**D**). Data shown as the mean value of 6 or more individual replicates, with error bars indicating SD. Filled symbols represent animals with placental transmission, and non-filled symbols represent those without transmission. (**E**) Compared to the control group, median peak plasma viral load (VL) is reduced by nearly 2 logs in high-potency HIG group (p=0.047, corrected Wilcoxon exact test). (**F**) There is a correlation between peak maternal plasma VL and initial amniotic fluid VL (r=0.812, p=0.002, nonparametric Spearman correlation). Additionally in comparison to the control group, the average number of days to first urine shedding (**G**) and first saliva shedding (**H**) is increased in the high-potency HIG group (both p=0.047, corrected Wilcoxon exact test). Horizontal bars indicate median values. *p<0.05, corrected Wilcoxon exact test.

### Short NGS Amplicon Population Profiling (SNAPP) of plasma virus

Using a novel, validated (Fig. S6) next generation sequencing technique, short 400bp regions within the RhCMV gB (Fig. 5) and gL (Fig. S7) genetic loci were amplified and sequenced at great read depth. We found that, as previously reported,^24^ the viral populations in plasma were composed primarily of strain UCD52 at each sampled time point and in all treatment groups. Additionally, viral diversity was measured by calculating mean nucleotide diversity (*π*) across each sample’s haplotype sequences (Table S2). Overall, the diversity of the dominant plasma UCD52 viral subpopulation was restricted in the high-potency HIG-treated animals compared to the control group at both the gB (Fig. 5C) and gL (Fig. S7) loci (both p=0.036, Wilcoxon exact test). Nucleotide diversity was not significantly different between the control and standard HIG groups.

**Fig. 5.**
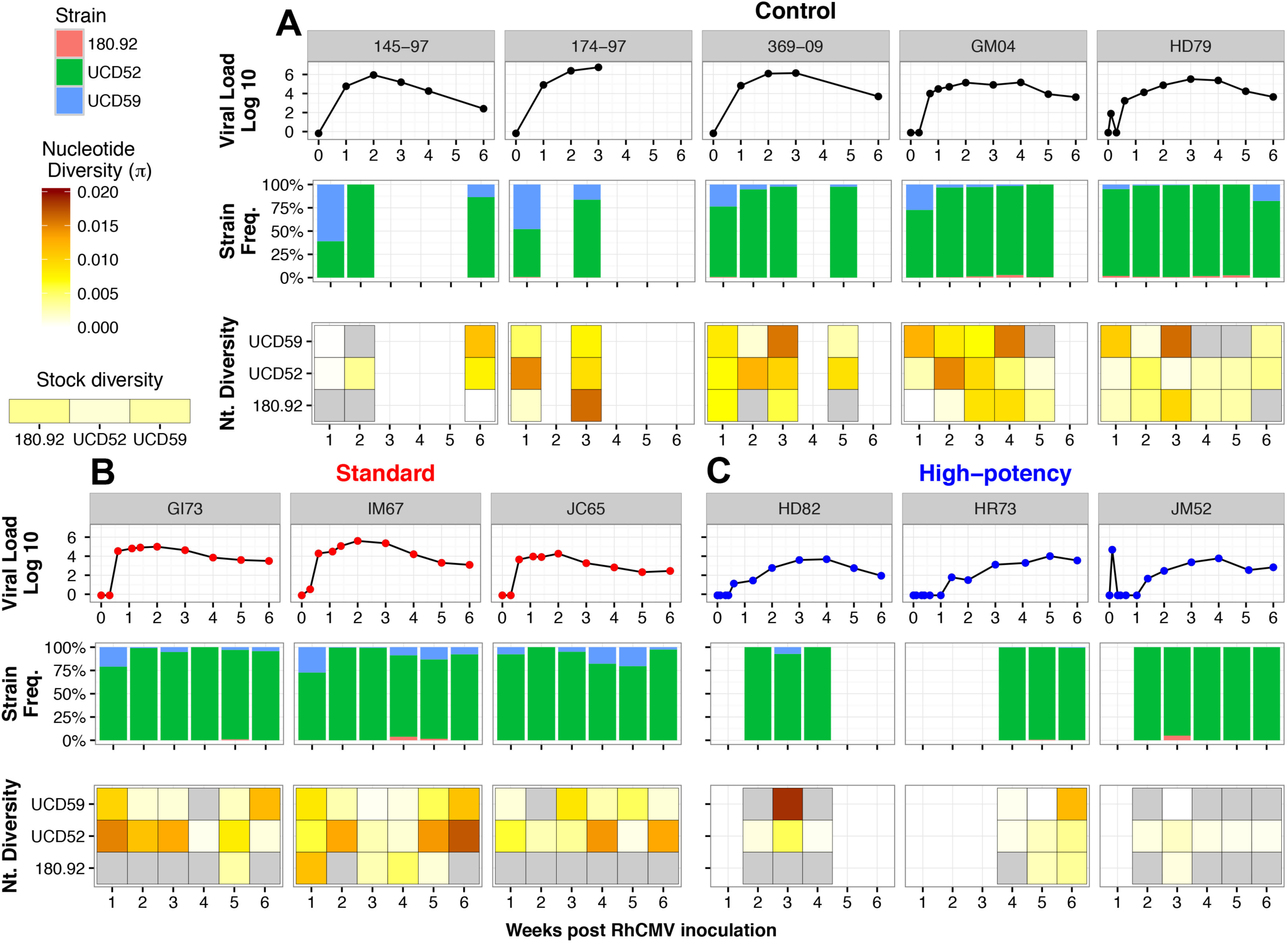
Plasma RhCMV gB sequence diversity is decreased following high-potency HIG infusion. SNAPP deep sequencing analysis at the glycoprotein B locus for maternal plasma virus population is shown for (**A**) 5 control dams (145-97, 174,97, 369-09, GM04, and HD79), (**B**) 3 standard HIG infused dams (GI73, IM67, and JC65), and (**C**) 3 high-potency HIG infused dams (HD82, HR73, and JM52). The **top panel** for each animal indicates the plasma viral load. The **middle panel** indicates the percentage of sequence reads corresponding to each of the three inoculated RhCMV strains in plasma at weeks 1-6 post infection (180.92 in pink, UCD52 in green, and UCD59 in blue), which was similar between the treatment groups. The **bottom panel** for each animal is a heat map depiction of viral nucleotide diversity (π). Scale ranges from white (π=0) to dark red (π=0.02), with the diversity of each viral stock displayed for reference. Gray coloring indicates that no sequence reads were detected at that time point for a given viral variant. Blank areas represent sample non-availability (control group) or limited plasma viral load (<100 copies/mL). There is a significant reduction in diversity at the gB locus for the dominant strain in plasma (UCD52) following high-potency HIG infusion (*p=0.036, Mann-Whitney U test).

### Fetal outcome and RhCMV congenital infection

While 5 of 6 control dams aborted their fetus at 3 weeks post RhCMV infection, administration of either HIG regimen was sufficient to protect fetuses from spontaneous abortion (p=0.015, exact log-rank test based on Heinze Macro; Fig. 6A).^30^ Furthermore, we observed a reduced rate of congenital RhCMV transmission among HIG-treated animals (2/6) compared to control animals (6/6) (p=0.049, exact log-rank test based on Heinze Macro; Fig. 6B).^30^ Intriguingly, in the standard HIG group only 1 of 3 fetuses remained uninfected whereas all 3 fetuses in the high-potency HIG group were protected from infection (Fig. 6C), suggesting that preexisting, potently CMV-neutralizing HIG can reduce the incidence of congenital infection. Of note, fetal growth curves (Fig. S8) and systemic cytokine profiles (Fig. S9) were similar between treatment groups, indicating no obvious off-target effects of the infused HIG. Reassuringly, all dams with detectable RhCMV DNA in amniotic fluid also had detectable virus in placenta, amniotic membrane, and umbilical cord (Fig. 6C). However, there was significant heterogeneity in viral burden and distribution among fetal tissue types: some fetuses had high copy number virus in nearly every tissue tested (174-97, IM67), while others only had virus detectable in the cochlea (GI73, HD79) (Fig. 6C). The persistence of virus in the cochlea further validates this model as consistent with congenital HCMV infection in human fetuses. Congenital infection was additionally confirmed in placentas by immunohistochemical staining for the RhCMV IE-1 protein (Fig. 6D, Table S3).

**Fig. 6.**
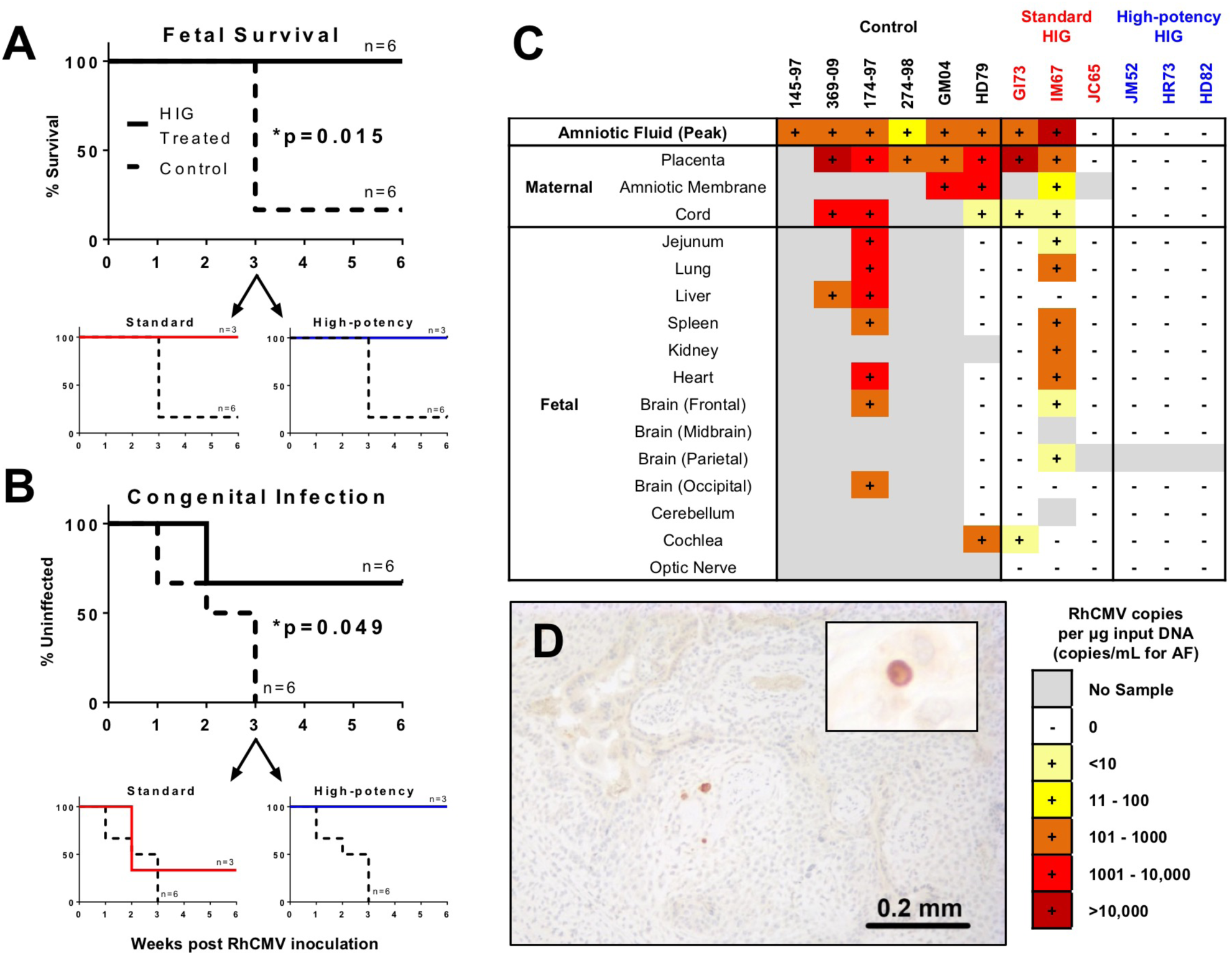
Maternal HIG pre-infusion significantly reduces incidence and severity of RhCMV congenital infection. (**A**) Fetal survival was significantly increased following HIG infusion (p=0.015, Exact log-rank test based on Heinze Macro), and both standard HIG and high-potency HIG infusion were sufficient to prevent fetal loss. (**B**) Rate of congenital infection was significantly reduced following HIG pre-infusion (p=0.049, Exact log-rank test based on Heinze Macro), though 2/3 animals in the standard HIG group had congenitally-infected infants compared with 0/3 in the high-potency HIG group**. (C**) Heat map separated by treatment group demonstrates heterogeneity in the viral burden of amniotic fluid, placenta, amniotic membrane, cord, and fetal tissues. (**D**) Placental infection was detected by immunohistochemistry for RhCMV IE-1 protein, with rare cells in the trophoblastic shell of IM67 (Standard HIG) exhibiting intranuclear staining. Inset: higher magnification of cell exhibiting intranuclear staining.

### Placental transcriptome

Congenital RhCMV infection appears to modify the placental transcriptome more radically than HIG infusion or fetal abortion, as more than 300 genes were differentially-expressed between RhCMV-transmitting and nontransmitting dams, but only 78 when comparing HIG infusion/fetal outcome (Fig. 7A, Fig. S10). This finding suggests that HIG predominantly mediated protection through its impact on maternal systemic virus replication rather than through modification of the placental transcriptome. A large proportion of these differentially-expressed genes were immune-related. Additional cellular functions that appeared altered by RhCMV infection included metabolism, oxidoreductase activity, and cellular growth/proliferation (Table S4). Gene interaction analysis of differentially-expressed genes between RhCMV-transmitting and nontransmitting dams (Fig. 7B) revealed that the placental transcriptome was heavily biased towards up-regulation of genes in transmission. Furthermore, certain gene nodes had numerous connections to other differentially regulated genes (e.g. VCAM1, EGFR, GZMB, LYN, FYN, GAPDH), suggesting that these “keystone” genes are central to the cellular pathways induced by RhCMV infection. Focusing on a subset of genes involved in innate vs. adaptive immunity (Fig. 7C), we note that there is a potential bias for activation of innate immune pathways by placental RhCMV infection. Finally, cell type enrichment analysis based upon previously-described genes involved in cellular activation/processes^31^ demonstrated no preference at the transcriptome level for myeloid vs. lymphoid lineages (Fig. 7D). However, Natural Killer (NK) cell-specific genes had the most robust differential expression (Fig. 7D), and we identified 21 unique genes known to be associated with NK cell function and/or chemotaxis (Fig. 7E, Table S5) including killer immunoglobulin-like receptors (KIR2DL4) and killer lectin-like receptors (KLRD1, KLRC1, KLRC3, KLRB1). Furthermore, non-classical MHC molecules HLA-E and HLA-G (rhesus analog inferred by IPA database) were up-regulated.

**Fig. 7.**
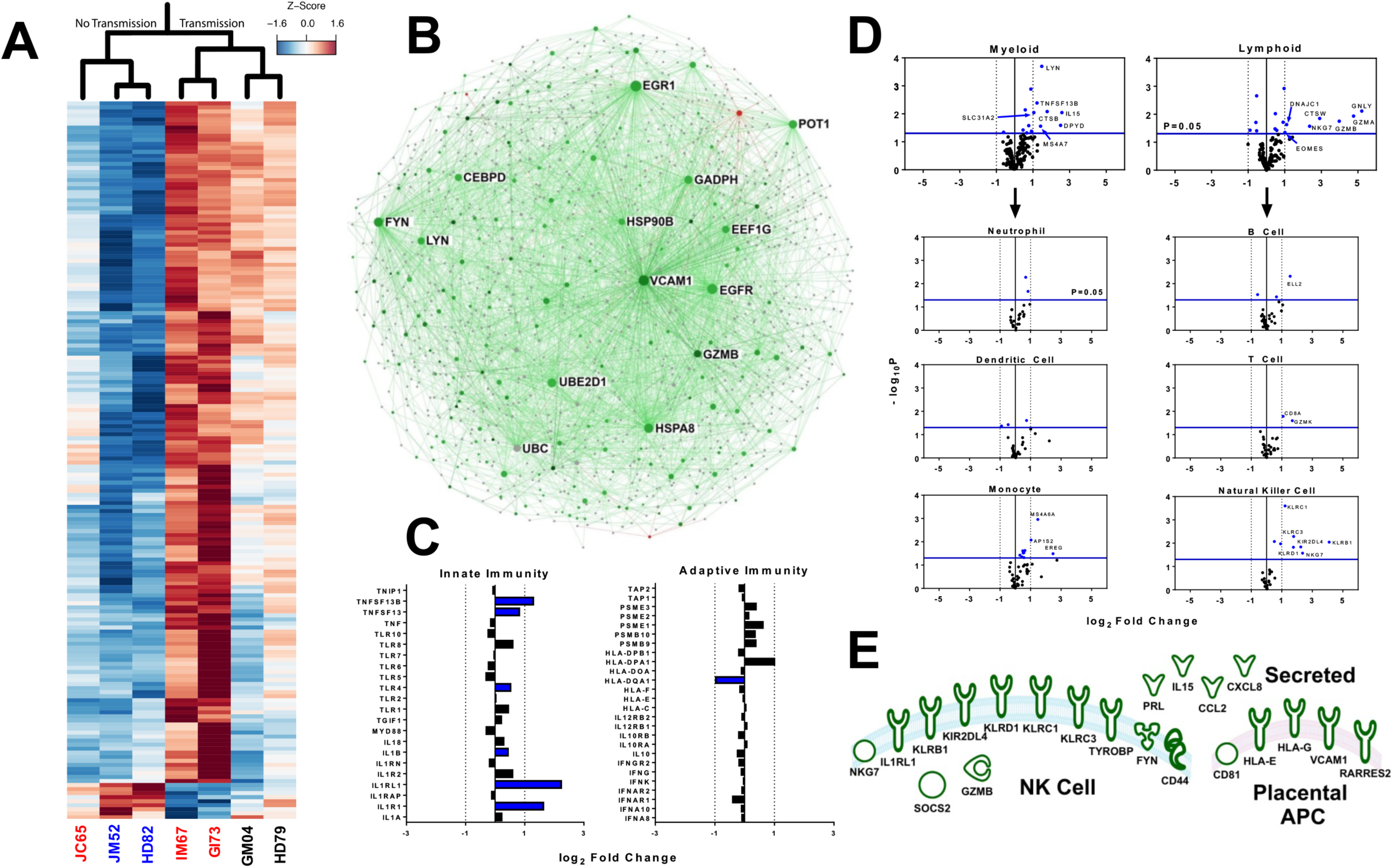
Placental transcriptome shifts following RhCMV congenital infection. (**A**) Heat map depicting normalized fold change of top differentially expressed genes (p<0.01, fc>2.0) in monkey dam placenta with and without RhCMV congenital infection (red=relative up-regulation in RhCMV infection, blue=relative down-regulation) (**B**) Interaction network for all differentially regulated genes (p<0.05, fc>2.0) suggests that genes up-regulated in congenital infection (green) greatly outnumber those up-regulated in no infection (red). Node size reflects the number of gene interactions (InnateDB database), while color intensity indicates degree of fold change. (**C**) Analysis of innate vs. adaptive immune genes suggests some preference for up-regulation of innate immunity following infection, with blue denoting a statistically significant change (p<0.05) (**D**) Cell type enrichment analysis suggests no preference for myeloid vs. lymphoid lineages, but that genes specific for monocytes and NK cells are up-regulated following congenital infection. Statistically significant genes (p<0.05) shown in blue, and significant genes with a fc>2.0 are labeled. (**E**) Diagram depicting placental genes related to NK cell movement and/or function (defined by IPA and KEGG databases) that are differentially up-regulated (p<0.05, fc>2) in placental RhCMV infection.

## Discussion

The role of preexisting antibodies in prevention of congenital HCMV infection remains controversial within the HCMV vaccine field, yet our findings indicate that a dose-optimized preinfusion with potently-neutralizing HIG can block placental RhCMV transmission in a rhesus monkey model of congenital CMV. Our study illustrates that factors such as maternal antibody titer and neutralization potency at the time of peak CMV viremia may be important considerations for antibody-mediated protection against congenital CMV infection. In this study, given the experimental setup, we have not definitively proven that it is neutralization titer (and not other factors such as total amount of binding antibody, antibody-dependent cellular cytotoxicity, or reduction in maternal viral load) that is protective against congenital RhCMV transmission. Nevertheless, these data substantiate observational human cohort studies suggesting that maternal HCMV-specific antibodies decrease risk of placental virus transmission.^20-22,32^ Furthermore, these data affirm findings from the guinea pig model of congenital infection that demonstrate that preexisting antibodies (from either glycoprotein immunization or passive HIG/monoclonal antibody infusion) can reduce the incidence and/or severity of congenital gpCMV infection.^33-36^

The therapeutic efficacy of HIG in preventing placental transmission in mothers with primary HCMV infection is a topic of ongoing investigation. Several non-randomized small-scale clinical trials^37,38^ and a case-control study^39^ concluded that post-infection HIG infusion significantly reduced rates of placental HCMV transmission and improved infant outcomes. Animal studies in the guinea pig model of congenital infection further corroborated these findings.^35,40^ However, it was perhaps unsurprising when a large-scale, randomized, placebo-controlled trial did not find significant reduction in rates of congenital infection in mothers with primary HCMV infection that were subsequently treated with HIG, and revealed no difference in virus-specific antibody titers, effector T cell count, or the level of plasma viremia between the HIG-treated and placebo groups.^23^ Several confounding factors may explain the lack of efficacy of HIG treatment in this large-scale clinical trial. First, it was unknown in this study whether congenital transmission had already occurred prior to initiating therapeutic HIG treatment. Second, there is no concrete evidence for the therapeutic efficacy of the clinically-utilized product (Cytotect^®^), and therefore it is unknown whether HIG infusion achieved a titer that was both sustained and/or sufficient for effective anti-HCMV immunity.

This study was not intended to model a therapeutic treatment regimen but rather to directly address the clinical question: can preexisting antibodies alone prevent congenital CMV infection? The advantages of our NHP model are that the timing of infection is predefined and serum antibody titer can be titrated. By utilizing passive infusion experiments with different dosing regimens and potency of RhCMV-specific antibodies prior to RhCMV inoculation, we were able to model the requirements for preexisting antibodies to confer protection against congenital RhCMV infection. Moreover, conducting the experiments in a CD4+ T cell-depletion model also allowed us to address the protective role of antibodies to the exclusion of RhCMV-specific cellular immune responses. Our results suggest that a single 100mg/kg dose of standard neutralizing potency HIG (donor plasma screened by elevated ELISA OD, as is done for the clinical product Cytotect^®^ ^41^) delivered immediately prior to RhCMV infection can reduce the severity of congenital infection (0 of 3 monkeys aborted) but is insufficient to prevent congenital CMV transmission (2 of 3 monkeys transmitted). However, a dose-optimized preinfusion of a higher neutralizing potency HIG product may be more effective in preventing and/or decreasing the severity of congenital infection.

To our knowledge, the epitope specificity of RhCMV-neutralizing antibodies has not been previously reported. It has been well established for HCMV that, while gB is an important target of neutralizing antibodies, the majority of neutralization activity in seropositive individuals is directed against the gH/gL/UL128-131A pentameric complex.^42,43^ Furthermore, the pentameric complex has also been shown to be a target of neutralizing antibodies for guinea pig CMV.^44^ In this study, we identified that antibodies targeting both RhgB and RhPC contribute to the total neutralization activity of RhCMV-seropositive monkey sera in approximately equivalent proportions (Fig. 2). It is certainly reassuring for the translational applicability of this animal model system that RhPC-directed antibodies are neutralizing; however, it is unclear why the relative proportions of PC-directed neutralization activity would differ between HCMV (∼85%)^42^ and RhCMV (∼50%). We hypothesize that this could explain why the magnitude of RhCMV neutralizing antibody titers are roughly equivalent magnitude when measured in both fibroblasts and epithelial cells (Fig. 3), whereas HCMV neutralizing antibody titers are nearly an order of magnitude lower in fibroblasts than epithelial cells.^45^ Subsequent studies are needed to improve our understanding of this potential difference in viral biology and/or antiviral humoral immune factors between RhCMV and HCMV.

It has been theorized that passively-infused antibodies can reduce systemic maternal viral load and thereby reduce the likelihood of placental transmission,^20,37^ though this correlation has not been observed in some human primary infection cohorts.^46^ In this study, dams with reduced peak plasma viral load in the presence of antibodies did not transmit the infection to their fetus *in utero* (Fig. 4E). This trend was accompanied by a significant reduction in nucleotide diversity of the plasma virus population in the high-potency HIG group, assessed at both the gB and gL loci (Fig. 5, Fig. S7). Thus, we hypothesize that in the setting of antibody-mediated immune pressure, the number of unique viral variants is restricted following primary infection and systemic replication is inhibited, resulting in decreased viral transmission to the fetus.

The placental transcriptome was examined to investigate whether HIG-mediated protection against congenital infection occurs via gene regulation at the level of the placenta/decidua. Intriguingly, RhCMV infection impacted the transcription profile of placental/decidual cells to a much greater degree than HIG-treatment or fetal abortion, suggesting that HIG predominantly mediated protection through its impact on maternal systemic virus replication. The preferential activation of innate over adaptive immune processes in RhCMV-infected placenta (Fig. 7D) is perhaps due to the tolerogenic environment at the maternofetal interface.^47,48^ Furthermore, the apparent upregulation of NK cell-specific genes in RhCMV-infected placenta (Fig. 7D,E) may be attributable to either NK cells being the major leukocyte in the decidua during the early stages of pregnancy^49^ or by the propensity for NK cells to control CMV infection *in vivo*.^50,51^

Our findings are based on a relatively small cohort of animals (n=12) due to the severely limited availability of breeding, seronegative female monkeys. One consequence of our limited animal population is the need to amplify the biologic effects of placental RhCMV infection through both CD4+ T cell depletion and IV viral inoculation. Though both these artificial methods distort the reality of natural CMV infection and bias our results in favor of placental transmission, we propose that these interventions strengthen the major finding in this study that preexisting antibody alone (in the absence of cell-mediated immunity) can prevent placental transmission in the setting of acute maternal viremia. One additional limitation to our study design is that, though HIG is known to non-specifically modulate immune responses to prevent inflammatory disease,^52-54^ we had to limit the number of individual control groups in this study and did not have a cohort of animals infused with non RhCMV-reactive HIG. Because of this deficiency, we cannot definitively rule out non-specific immunomodulatory effects of the infused HIG, though all infusion groups had similar cytokine expression profiles following infection (Fig. S9). Furthermore, since protection against congenital transmission may have been achieved as a result of a “dose-effect” in the high-potency HIG group vs. the standard group, antibody-mediated protection is likely due to anti-viral function rather than immunomodulatory properties of the infused HIG.

Despite these limitations, this is the first primate study to demonstrate that preexisting maternal antibody alone has the potential to block and ameliorate placental CMV transmission. Thus, a maternal vaccine that elicits durable, potently-neutralizing humoral immunity could be an ideal intervention to eliminate pediatric disabilities associated with congenital HCMV infection. Based on our findings we hypothesize that the prevention of maternal HCMV acquisition (i.e. sterilizing immunity), which has been the target of several phase II vaccine trials,^13,14^ may not be the most appropriate measurable outcome for a congenital HCMV vaccine that ultimately seeks a reduction in the frequency of congenital HCMV transmission. Since we observed that antibody-mediated protection against congenital HCMV was associated with a decrease in maternal systemic viral load and constrained viral diversity, these surrogate markers could potentially serve as clinical study endpoints. Subsequent studies should investigate the epitope specificity and properties of protective immunoglobulins for future rational design of a maternal congenital HCMV vaccine.

## Methods

### Animals

Indian-origin rhesus macaques were housed at the Tulane Primate National Research Center and maintained in accordance with institutional and federal guidelines for the care and use of laboratory animals.^55^ All females were from the expanded specific pathogen-free (eSPF) colony and confirmed to be RhCMV seronegative by whole-viron ELISA screening for RhCMV-specific IgM and IgG plasma antibodies. RhCMV-seronegative males and females were then co-housed and females were screened every 3 weeks for pregnancy via abdominal ultrasound. All animals in this study were between 4 and 9 years of age at the time of pregnancy/enrollment (JC65=4.7y, GI73=8.8y, IM67=5.7, JM52=4.6, HR73=7.6, HD82=8.7, HD79=8.7, GM04=8.8, HD79=8.7). Gestational dating was performed by sonography, and was based on gestational sac size (average of three dimensions) and crown rump length of the fetus. At week 7 of gestation, 12 RhCMV-seronegative dams were administered an IV 50mg/kg dose of recombinant rhesus CD4+ T cell–depleting antibody (CD4R1 clone; Nonhuman Primate Reagent Resource).

One week following CD4+ T cell depletion, 3 seronegative dams were administered a single dose (100mg/kg) of hyperimmune globulin (HIG) produced from the plasma of RhCMV seropositive monkeys screened by RhCMV whole virion IgG binding ELISA (50% Inhibitory Concentration-IC_50_=22.64μg/mL) 1hr prior to intravenous (IV) challenge with a mixture of RhCMV strains (2×10^6^ TCID50 RhCMV 180.92^27^ in one arm and 1×10^6^ TCID_50_ each of RhCMV UCD52 and UCD59^28^ in the other arm, diluted in serum-free RPMI). The remaining 3 dams received a dose-optimized (2 dose) regimen of a potently-neutralizing HIG product purified from RhCMV seropositive monkeys with a plasma epithelial cell neutralization titer >1000 (IC_50_=5.88μg/mL) at 1hr prior to IV RhCMV inoculation (150mg/kg) and 3 days post infection (100mg/kg), respectively. This dose-optimized regimen was based on pharmacokinetic modeling of the neutralization activity from the standard group (Fig. S4). 6 dams were non-infused control animals (including 4 historical controls) ^24^.

Blood draws were performed on the standard potency infused animals on the following days after infection/infusion: 0, 2, 4, 7, 10, 14, 21, 28, 35, 42. The high-potency infused animals were sampled at days: 0, 1, 2, 3, 4, 7, 10, 14, 21, 28, 35, 42, 49, 56, 63, 70, 77, 84. Amniotic fluid (by amniocentesis), urine (by clean pan collection), and saliva (by oral saline wash), were also collected on all animals on a weekly to biweekly intervals until fetal harvest. In the case of fetal abortion, placental and fetal products were obtained from the cage and fixed in formalin for hematoxylin–eosin staining and immunohistochemistry or frozen. Fetal growth was monitored by measuring biparietal diameter and femur length during weekly to biweekly fetal sonography over the course of gestation. Sonography was also used to screen for signs of congenital RhCMV-associated sequelae. The fetuses of dams in the standard HIG group were harvested at 6 weeks following infection, and those of the high-potency HIG group at 12 weeks following infection.

### IgG purification for HIG preparations

Seropositive rhesus monkeys plasma donors were screened by either high OD binding against UCD52 virions (top 50% seropositive OD; standard HIG) or by epithelial cell neutralization titer (ID_50_>1000; high-potency HIG). Donor plasma was mixed with equal volume of Protein G IgG Binding Buffer (Pierce) and spun at 4700 RPM for 25 minute at RT. Pellet was discarded and supernatant was filter sterilized through a 0.22μm filter and used for purification. Sera mixture was passed through gravity column packed with Protein G Plus Agarose beads (Pierce). Beads were washed with 5 column volume (CV) of IgG Binding buffer and eluted with 2 CV of IgG Elution Buffer (Pierce). Eluted IgG was immediately neutralized with 1M Tris buffer, pH=8. Beads were washed with 5 CV of IgG binding buffer. Serum was passed through Protein G Plus Agarose beads 10 times and IgG was eluted each time. All IgG elutions were concentrated and buffer exchanged into 1x PBS using Millipore Amicon 30K filters (Fisher Scientific). Final preparations were tested by ELISA for the presence of bacterial endotoxin. See quality control sheet for high-potency HIG preparation in Fig. S3.

### Pharmacokinetic modeling for dose-optimized HIG infusion

Plasma concentration-time data of IgG in rhesus monkeys were adjusted using baseline value of IgG for each individual animal and analyzed using WinNonlin (version 6.3.0; Certara). One- and two-compartment PK models with first-order elimination were evaluated to characterize the observed data. PK parameters including clearance (CL), volume of distribution (V), and half-life associated with the elimination phase (T_1/2,β_) were estimated for each animal. Simulations for optimizing dosing regimen in the second PK study were performed to ensure IgG concentrations above 1-1.2 mg/ml (∼ID_50_>1000) for 2 weeks after initiating treatment.

### Cell culture and virus growth

Telomerized rhesus fibroblasts (TeloRF) cells were maintained in Dulbecco’s modified Eagle medium (DMEM) containing 10% FCS, 2mM L-glutamine, 50U/mL penicillin, 50μg/mL each streptomycin and gentamicin, and 100μg/mL geneticin G418 (Invitrogen). Monkey kidney epithelial (MKE) cells were cultured in DMEM-F12 supplemented with 10% FCS, 2mM L-glutamine, 1mM sodium pyruvate, 50U/mL penicillin, 50μg/mL each streptomycin and gentamicin, and 5mL of epithelial growth cell supplement (ScienCell). Cell lines were tested for mycoplasma contamination every 6 months.

Epithelial-tropic UCD52 and UCD59 were propagated on MKE cells for 3-4 passages, then passaged once on TeloRF cells in order to increase viral titer. This single round of fibroblast amplification was intended to minimize any alteration of viral tropism or glycoprotein surface expression. Virus infections were performed with similar media but with 5% FCS for 4h at 37°C and 5% CO_2_. Virus was harvested when cells showed 90% cytopathic effects (CPE) by cell scraping. For harvest, infected cells were pelleted by low-speed centrifugation, and the supernatant was placed on ice. Cell pellets were then resuspended in infection media and subjected to three rounds of freeze/thaw cycles. Following centrifugation, supernatants were combined, passed through a 0.45μm filter, overlaid onto a 20% sucrose cushion, and ultracentrifuged at 70,000×*g* for 2h at 4 °C using an SW28 Beckman Coulter rotor. Virus pellets were resuspended in DMEM containing 10% FCS and titered in TeloRFs using the TCID_50_ method of Reed and Muench.

### Characterization of T cells by flow cytometry

Maternal peripheral CD4+ and CD8+ T cells were phenotyped by mixing 100μL of EDTA-anticoagulated blood with a pool of fluorescently labeled monoclonal antibodies including CCR7, CD95, CD28, CD4, CD20, CD3, CD8, and CD45 (Table S1). After a 30 min incubation at 4°C, cells were washed with PBS supplemented with 2% FCS and pelleted at 100×*g* for 5 min. Red blood cells were then lysed by adding 1×BD lysis buffer for 40min at room temperature. Intact cells were centrifuged at 150×*g* for 5 min, washed once with PBS containing 2% FCS, and resuspended in 100μL of PBS with 2% FCS. Cells were fixed by adding 15μL of 2% formaldehyde and processed by flow cytometry. Gating of CD4+ and CD8+ populations/subpopulations was completed in FlowJo as is outlined in Fig. S1.

### Antibody kinetics and glycoprotein binding specificity

Virus-specific IgM and IgG antibody kinetics were measured in maternal plasma by whole-virion ELISAs, and glycoprotein specificity of HIG and maternal plasma samples was measured by ELISA against soluble RhCMV gB protein and RhCMV pentameric complex. Plates were coated with antigen diluted in PBS with Mg^2+^ and Ca^2+^ (PBS+) overnight at 4°C – either 5,120 PFU/mL of filtered RhCMV UCD52 virus for whole virion ELISA or 0.13μg/mL soluble glycoprotein. Following incubation, plates were blocked for 2h with blocking solution (PBS+, 4% whey protein, 15% goat serum, 0.5% Tween-20), then threefold serial dilutions of plasma (1:30 to 1:5,314,410) were added to the wells in duplicate for 1.5h. Plates were then washed twice and incubated for 1hr with anti-monkey IgM or anti-monkey IgG HRP-conjugated antibodies (Rockland) at a 1:10,000 dilution. After four washes, SureBlue Reserve TMB Microwell Peroxidase Substrate (KPL) was added to the wells for 10min, and the reaction was stopped by addition of 1% HCl solution. Plates were read at 450nm. The lower threshold for antibody reactivity was considered to be 3 SDs above the average optical density measured for the preinfection, RhCMV seronegative samples at the starting plasma dilution (1:30).

### Neutralization assays

TeloRF and MKE cells were seeded into 96-well plates and incubated for 2 days at 37°C and 5% CO_2_ to achieve 100% confluency. After 2 days, serial dilutions (1:10 to 1:30,000) of heat-inactivated rhesus plasma were incubated with RhCMV 180.92 or RhCMV UCD52 (MOI=1) in a 50μl volume for 45min at 37°C. The virus/plasma dilutions were then added in duplicate to wells containing TeloRF or MKE cells, respectively, and incubated at 37°C for 2h. After washing, cells were incubated at 37°C for an additional 16h. Infected cells were then fixed for 20min at -20°C with 1:1 methanol/acetone, rehydrated in PBS with Mg^2+^ and Ca^2+^ (PBS+) 3×5min, and processed for immunofluorescence with 1μg/mL mouse anti-RhCMV IE-1 monoclonal antibody (provided by Dr. Daniel Cawley, Oregon Health and Science University) followed by a 1:500 dilution of goat anti-mouse IgG-Alexa Fluor 488 antibody (Millipore). Nuclei were stained with DAPI for 5min (Pierce). Infection was quantified in each well by automated cell counting software using either: (1) a Nikon Eclipse TE2000-E fluorescent microscope equipped with a CoolSNAP HQ-2 camera at 10× magnification OR (2) a Cellomics Arrayscan VTI HCS instrument at 10× magnification. Subsequently, the 50% inhibitory dose was calculated as the sample dilution that caused a 50% reduction in the number of infected cells compared with wells treated with virus only using the method of Reed and Muench.

### Soluble protein bead coupling and antibody depletion

Cyanogen bromide (CNBr) activated Sepharose beads (GE Healthcare) were re-hydrated in 1mM HCl, then suspended in coupling buffer (0.1M NaHCO_3_ + 0.5M NaCl, pH 8.3) at room temperature (RT). Every 100μL of bead slurry was combined with 150μg soluble RhgB or RhPC ligands. Coupling was allowed to proceed for 1h at RT on an inversion rotator. Excess soluble protein was washed off with 5 column volumes coupling buffer. CNBr unbound active groups were blocked by incubation in 0.1M Tris HCl, pH 8.0 for 2h at RT on inversion rotator. Protein-bound beads were washed with 3 cycles of alternating pH: first 0.1M acetic acid, pH 4.0, then second 0.1M Tris-HCl, pH 8.0. Beads were suspended in PBS and stored at 4°C. For depletion of antibodies for a given specificity, 100μL of protein-coupled bead slurry were loaded into a spin microelution column (Pierce, TFS). 50μL of plasma was filtered (0.22μM), then loaded into the column. Plasma was centrifuged through the column ×5 without elution between steps. Bound IgG was eluted using a 0.2M glycine elution buffer, pH 2.5. Column was recalibrated by washing with 3 cycles of solutions of alternating pH (as described above). Adequate and specific depletion of plasma was confirmed by ELISA against the depleted protein (>75% change in EC_50_ against depleted epitope; <20% change in EC_50_ against non-depleted epitope).

### Tissue processing and staining

Standard immunoperoxidase staining for RhCMV was performed on formalin-fixed, paraffin-embedded sections of multiple tissues. Tissue sections were deparaffinized in xylene, rehydrated in graded alcohol, and subsequently blocked with 0.3% hydrogen peroxide in PBS for 30min. Pretreatment involved microwaving for 20min in 0.01% citrate buffer (Vector laboratories), followed by 45min of cooling at room temperature. Following pretreatment, an avidin–biotin block (Invitrogen) and protein block with 10% normal goat serum (NGS) were conducted on all sections. A wash of Tris-buffered saline with 0.5% Tween-20 followed each step. Sections were incubated with anti–RhCMV IE1 polyclonal rabbit sera (kindly provided by P.A.B.) at a 1:1,600 dilution for 30 min at room temperature. Slides were then incubated with biotinylated goat anti-rabbit IgG (Invitrogen) at a 1:200 dilution in NGS for 30min at room temperature. This was followed by 30min incubation at room temperature with R.T.U. Vectastain ABC (Vector Laboratories). Immunolabeling was visualized using diaminobenzidine and counterstained with Mayer’s hematoxylin. Irrelevant, isotype-matched primary antibodies were used in place of the test antibody as negative controls in all immunohistochemical studies. Positive control tissues consisted of archived rhesus macaque lung and testis from an RhCMV-seropositive animal.

### Viral DNA isolation and assessment of viral load

RhCMV viral load was measured in plasma, AF, saliva, urine, and tissues by quantitative RhCMV-specific PCR following DNA extraction. Urine and mouth washes were concentrated using Ultracel YM-100 filters (Amicon) and frozen for subsequent DNA extraction. DNA was extracted from urine using the QIAmp RNA minikit (Qiagen), from AF/saliva/plasma by using the QIAmp DNA minikit, and from snap-frozen tissue (10-25mg) after overnight Proteinase K digestion using the DNeasy Blood and Tissue kit (Qiagen). RhCMV DNA was quantitated by real-time PCR using the 5′- GTTTAGGGAACCGCCATTCTG-3′ forward primer, 5′-GTATCCGCGTTCCAATGCA-3′ reverse primer, and 5′-FAM-TCCAGCCTCCATAGCCGGGAAGG-TAMRA-3′ probe, which amplify and detect a 108-bp region of the RhCMV IE1 gene.^56^ Between 5-10ng of DNA was added as template to 50μL of 1×PCR mixture containing 300nM of each primer, 100nM probe, 2mM MgCl_2_, 200μM each of dATP, dCTP, and dGTP, 400μM dUTP, 0.01U/μL Amperase UNG, 0.025U/μL Taq polymerase, and reaction buffer that includes passive reference dye ROX. PCR conditions consisted of an initial 2min cycle at 50°C followed by 10min at 95°C and 45 cycles of denaturation at 95°C for 15s and combined annealing/extension at 60°C for 1min. Data are expressed as copies per milliliter for plasma/AF and as copies per microgram input DNA for tissues/saliva/urine. RhCMV DNA was quantitated using a standard consisting of a plasmid containing the entire RhCMV IE1 gene. RhCMV DNA was detected with a linear dynamic range from 10^0^ to 10^6^ copies in the presence of genomic DNA.

### SNAPP (Short NGS Amplicon Population Profiling)

Viral DNA was isolated from plasma using the High Pure Viral Nucleic Acid kit (Roche Life Science). Hypervariable regions 400bp in length within glycoprotein B and glycoprotein L were amplified in duplicate by a single round of PCR using 5′-AAGTGCTCGAAGGGCTTCTC-3′ and 5′-TCTGGACATTGATCCGCTGG-3′ for the gB region and 5′-GCGCGGCACACATTATCTAC-3′ and 5′-GGTGAGTGCTGCTGTTTTGG-3′ for the gL region. Overhang regions were conjugated to primer locus-specific sequences for subsequent Illumina index primer addition and sequencing: forward primer overhang= 5′-TCGTCGGCAGCGTCAGATGTGTATAAGAGACAG-[locus]-3′ and reverse primer overhang= 5′-GTCTCGTGGGCTCGGAGATGTGTATAAGAGACAG-[locus]-3′. Approximately 10ng DNA was added as template to 50μL of 1×PCR mixture containing 100nM of each primer, 2mM MgCl_2_, 200μM each of dNTP mix (Qiagen), 0.025U/μL Phusion Taq polymerase. PCR conditions consisted of an initial 2min denaturation at 98°C, followed by the minimum number of PCR cycles to achieve adequate amplification (98°C for 10s, 65°C for 30s, and 72°C for 30s), and a final 72°C extension for 10min. Products were purified using Agencourt AMPure XP beads (Beckman Coulter), then Illumina Nextera XT index primers were added by 15 cycles of amplification. The resultant PCR product was gel purified using ZR-96 Zymoclean Gel DNA Recovery Kit (Zymogen). The molar amount of each sample was normalized by qPCR using the KAPA library amplification kit (KAPA Biosystems). The library of individual amplicons was pooled together, diluted to an end concentration of 10pm, combined with 20% V3 PhiX (Illumina), then sequenced on Illumina Miseq using a 600 cycle V3 cartridge (Illumina). For each primer set we confirmed that there was no significant primer bias through mixing viral DNA in known ratios and applying the SNAPP technique (Fig. S6).

### SNAPP amplicon reconstruction and nucleotide diversity

Sequences of the targeted regions on the genes gB and gL were reconstructed by merging the paired reads sets using the PEAR software under default parameters.^57^The fused reads were then filtered using the extractor tool from the *SeekDeep* pipeline (http://baileylab.umassmed.edu/seekdeep), filtering sequences according to their length, overall quality scores and presence of the primer sequences. To homogenize the number of sequences per sample due to variable coverage, a random set of at most 10,000 sequences was retrieved from each sample to estimate strain frequency and nucleotide diversity. The number of sequences analyzed per sample varied from 2,000-10,000, depending on the sample coverage. Next, haplotype reconstruction was performed using the *qluster* tool from *SeekDeep*, which accounts for possible sequencing errors by collapsing fragments with mismatches at low quality positions. For each given sample, the haplotypes had to be present in two sample replicates to be taken into account. Each haplotype was assigned to 1 of the 3 inoculated viral strains by first calculating the nucleotide distance (nucleotide substitutions) between the haplotype and the reference viral strain, then assigning the haplotype to the strain with the shortest nucleotide distance. Nucleotide diversity was computed as the average distance between each possible pair of sequences belonging to the same reference strain.^58^

### Placental RNA isolation and microarray

Tissue sections weighing approximately 30mg were obtained from flash frozen, full-thickness rhesus placenta/decidua, lysed using a TissueLyser LT (Qiagen), then total RNA extracted using an RNeasy Mini Kit (Qiagen). RNA was assessed for quality with Agilent 2100 Bioanalyzer G2939A (Agilent Technologies) and Nanodrop 8000 spectrophotometer (Thermo Scientific/Nanodrop) and confirmed to have: A_260_/A_280_>1.8, A_260_/A_230_>1.0, and RIN>7.0. Hybridization targets were prepared with MessageAmp^™^ Premier RNA Amplification Kit (Applied Biosystems/Ambion) from total RNA, hybridized to Affymetrix GeneChip^®^ Rhesus Macaque Genome arrays in Affymetrix GeneChip^®^ hybridization oven 645, washed in Affymetrix GeneChip^®^ Fluidics Station 450 and scanned with Affymetrix GeneChip^®^ Scanner 7G according to standard Affymetrix GeneChip^®^ Hybridization, Wash, and Stain protocols (Affymetrix). Affymetrix microarray data was initially processed, underwent QC, and was normalized with the robust multi-array average method using the *affy^59^* Bioconductor^60^ package from the R statistical programming environment (R studio version 0.99.902). Differential expression was carried out using a moderated t-statistic from the *limma^61^* package. The false discovery rate was used to control for multiple hypothesis testing. Gene set enrichment analysis^62^ was performed to identify differentially regulated pathways and gene ontology terms for each of the comparisons performed. Gene sets specific for innate/adaptive immunity, myeloid/lymphoid lineages, and immune cell specific were obtained from previous reports^31^ and supplemented from the literature such that each comparison group had >25 characteristic genes (gene lists available upon request). Gene networks were constructed using NatureAnalyst^63^, with first order gene interactions inferred from the InnateDB interactome (Fig. 7B)^64^. Genes related to NK cell function (Table S5) were determined using QIAGEN’s Integrated Pathway Analysis (IPA) (NK cell activation and movement gene lists) and supplemented with those from the KEGG database (NK cell mediated cytotoxicity gene list).^65^

### Statistical analyses

Nonparametric tests were utilized because of the small study size. For viral load/shedding statistical analyses (Fig. 4E-H), Wilcoxon exact tests were Bonferroni-type corrected using the method of Benjamini and Hochberg to account for multiple comparisons.^66^ For sequence diversity analysis (Fig. 5, Fig. S7), UCD52 diversity was compared between treatment groups because it was the dominant strain replicating in plasma. First, median UCD52 nucleotide diversity was calculated for each animal. Median values were compared first by Kruskal-Wallis test (p=0.036), then by posthoc Mann-Whitney U test. Survival curve analyses were completed with an exact log-rank test based on Heinze macro.^30^ All statistical tests were carried out using the R statistical interface (version 3.3.1, www.r-project.org) and were two-tailed.

### Study approval

The animal protocol was approved by the Tulane University and the Duke University Medical Center Institutional Animal Care and Use Committees.

### Data and materials availability

SNAPP sequence data used in this manuscript is being compiled and will be made available in the NCBI SRA database prior to publication. Placental transcriptome microarray data is currently available in the NCBI GEO repository (GSE87395; http://www.ncbi.nlm.nih.gov/geo/query/acc.cgi?acc=GSE87395).

## Author Contributions

A.K. and S.R.P. designed research; C.S.N., D.T., K.M.B., M.G., R.B., X.A., and L.S. performed research; C.S.N., D.V.C., R.B., X.A., H.I., H.W., and M.C. analyzed data; A.D., F.C., F.W., D.J.D., N.V., M.W., P.A.B, M.C.W., K.K. contributed analytic tools/expertise/reagents; and C.S.N., A.K., and S.R.P. wrote the paper.

## Acknowledgements

The authors would like to recognize David Corcoran and the Duke Sequencing and Genomic Technologies Shared Resource for technical support, microarray data management, and feedback on the generation of the microarray data reported in this manuscript. Additionally, we would like to thank Dr. Xinzhen Yang and Pfizer Inc. for the generous gift of research materials. Finally, the excellent veterinary care and conduct of animal studies by the faculty and staff of the Departments of Veterinary Medicine and Collaborative Research at the Tulane National Primate Research Center (TNPRC) are gratefully acknowledged. The CD4+ T cell-depleting antibody used in these studies was provided by the NIH Nonhuman Primate Reagent Resource (R24 OD010976, and NIAID contract HHSN 272201300031C). This work was supported by: NIH/NICHD Director’s New Innovator grant to S.R.P (DP2HD075699), the TNPRC base grant NIH/NCRR (OD011104), NIH/NIAID grants to D.J.D. and P.A.B (R01AI103960, R01AI63356), and fellowship grant to C.S.N (F30HD089577). The funders had no role in study design, data collection and interpretation, decision to publish, or the preparation of this manuscript. The content is solely the responsibility of the authors and does not necessarily represent the official views of the National Institutes of Health.

